# Identification of a developmental switch in information transfer between whisker S1 and S2 cortex in mice

**DOI:** 10.1101/2021.08.06.455364

**Authors:** Linbi Cai, Jenq-Wei Yang, Shen-Ju Chou, Chia-Fang Wang, Heiko Luhmann, Theofanis Karayannis

## Abstract

The whiskers of rodents are a key sensory organ that provides critical tactile information for animal navigation and object exploration throughout life. Previous work has explored the developmental sensory-driven activation of the primary sensory cortex processing whisker information (wS1), also called barrel cortex. This body of work has shown that the barrel cortex is already activated by sensory stimuli during the first post-natal week. However, it is currently unknown when over the course of development these stimuli begin being processed by higher order cortical areas, such as secondary whisker somatosensory area (wS2). Here we investigate for the first time the developmental engagement of wS2 by sensory stimuli and the emergence of cortico-cortical communication from wS1 to wS2. Using in vivo wide-field imaging and electrophysiological recordings in control and conditional knock-out mice we find that wS1 and wS2 are able to process bottom-up information coming from the thalamus already right after birth. We identify that it is only at the end of the first post-natal week that wS1 begins to provide excitation into wS2, a connection which begins to acquire feed-forward inhibition characteristics after the second post-natal week. Therefore, we have uncovered a developmental window during which excitatory versus inhibitory functional connectivity between wS1 and wS2 takes place.

## 1. Introduction

Rodents are born with immature sensory systems (Khazipov et al. 2004; Leighton and Lohmann 2016). The postnatal development and maturation of brain circuits are essential for the detailed representation of environmental stimuli, which enables animals to interact with the external world in a refined manner (Ko et al. 2013). The somatosensory whisker system is key for rodent navigation and exploration of tactile stimuli already at birth (Akhmetshina et al. 2016; Yang et al. 2018). The information coming from the whiskers is conveyed via the trigeminal brainstem nuclei to the primary somatosensory thalamic nucleus, the ventro–posterior–medial nucleus (VPM), as well as the higher-order medial thalamic part of the posterior nucleus (POm). This information is then fed forward to the ‘barrels’ of the primary whisker somatosensory cortex (wS1), as well as the secondary whisker somatosensory cortex (wS2) and the motor cortex (Goebbels et al. 2006; Minlebaev et al. 2011; Staiger and Petersen 2021; Yamashita et al. 2013; Yamashita and Petersen 2016) . Anatomical studies have demonstrated dense reciprocal connections between wS1 and wS2 (Kwon et al. 2016; Minamisawa et al. 2018), and it has also been functionally shown that wS1 and wS2 represent whisker stimuli in a mirror-like topographic manner (Adibi 2019; Hubatz et al. 2020). In contrast to wS1, wS2 is preferentially activated by simultaneous whisker stimuli or single whisker stimuli delivered at high frequency (Menzel and Barth 2005). These findings, together with a shorter delay in arrival of sensory stimuli in wS1 compared to wS2 and larger receptive fields in wS2 vs. wS1, have suggested, as its name also indicates, that wS2 is a higher-order brain region that would process information coming from wS1. In this respect, the functional interaction between wS1 and wS2 has been studied in the behaving adult rodent, and coordinated activity from wS1 to wS2 has been shown to be essential for proper whisker-associated perception and learning behavior. For instance, wS2-projecting wS1 neurons show touch-related responses during a texture discrimination task (Chen et al. 2013) and they also develop specific patterns of activity during learning the task (Chen et al. 2015). Furthermore, wS2-projecting wS1 neurons have also been found to be involved in the goal-directed sensorimotor transformation of a whisker touch to licking motor output (Yamashita and Petersen 2016) and show higher choice-related activity than other neurons in layer 2/3 (Kwon et al. 2016). Despite of several studies about the development of sensorimotor processing (Dooley and Blumberg 2018; Gómez et al. 2021; Khazipov et al. 2004), little is known about the developmental stage that wS2 begins processing sensory information coming directly from the thalamus or indirectly via wS1, which would signify the beginning of higher-order representation of tactile stimuli in mammals. One informed hypothesis is that this occurs at the end of the second postnatal week. It is around postnatal day 14 when the animals begin to whisk and increase their locomotor activity searching for tactile information to help them navigate (Arakawa and Erzurumlu 2015; Landers and Philip Zeigler 2006).

Here, we have used a palette of *in vivo* approaches to assess the engagement of wS2 in whisker stimuli and probe its feed-forward activation by wS1 during development. By recording wS1 and wS2 activity *in vivo* simultaneously using either wide-field imaging or silicon probes, we find that wS2 processes sensory inputs from the first few days of postnatal life, activated directly through thalamocortical projections. It is only at the end of the first postnatal week that a long-lasting sensory-driven activity in wS1 starts propagating to wS2 driving its latent spiking phase, as shown through acute pharmacological manipulation of activity and genetic disruption of thalamocortical transmission. This excitatory drive between wS1 and wS2 dissipates by the end of the second postnatal week, which rather switches to an inhibitory one instead. Our work has identified a critical window of information transfer between wS1 and wS2, providing insights into the emergence of higher-order representation of the environment in the cortex.

## 2. Results

### The spatio-temporal development of sensory-evoked input activity in wS1 and wS2

Research in adult mice has shown that the wS2 displays a topographic map of the whisker pad, albeit more compact and less well defined compared to the wS1 (Hubatz et al. 2020). Here, we performed simultaneous transcranial wide-field voltage-sensitive dye imaging (VSDI) of wS1 and wS2, after deflecting individual whiskers of the contralateral whisker pad in four age groups across the first two weeks of postnatal life (P) (Fig. 1). In all age groups, single whisker deflection led to the clear activation of two centers in the first 300 ms after the stimulus (Fig. 1B), with wS1 preceding wS2 (Fig. 1C). We found that prior to P5, wS1 and wS2 displayed activity that remained local for the duration of the signal, but starting at P6, the activity expanded beyond the two discrete points and merged into a large cortical area covering both, suggesting that the interaction between wS1 and wS2 started at this time point. These results indicate that a rudimentary map of the whiskers is present in wS2 already within the first week after birth (Fig. 1A). For wS1 responses, a clear map of the whiskers is also already present at P3-4, which is in agreement with the characteristic segregation of thalamocortical fibers and the formation of the barrels around P3 (Erzurumlu and Gaspar 2012). These findings suggest that the thalamocortical fibers originating from the postero-medial (POm) thalamic nucleus or the ventral lateral part of the VPM (VPMvl), which project to wS2, are already functional early after birth and the functionally segregated columns largely appear at the end of the first week.

**Fig. 1.**
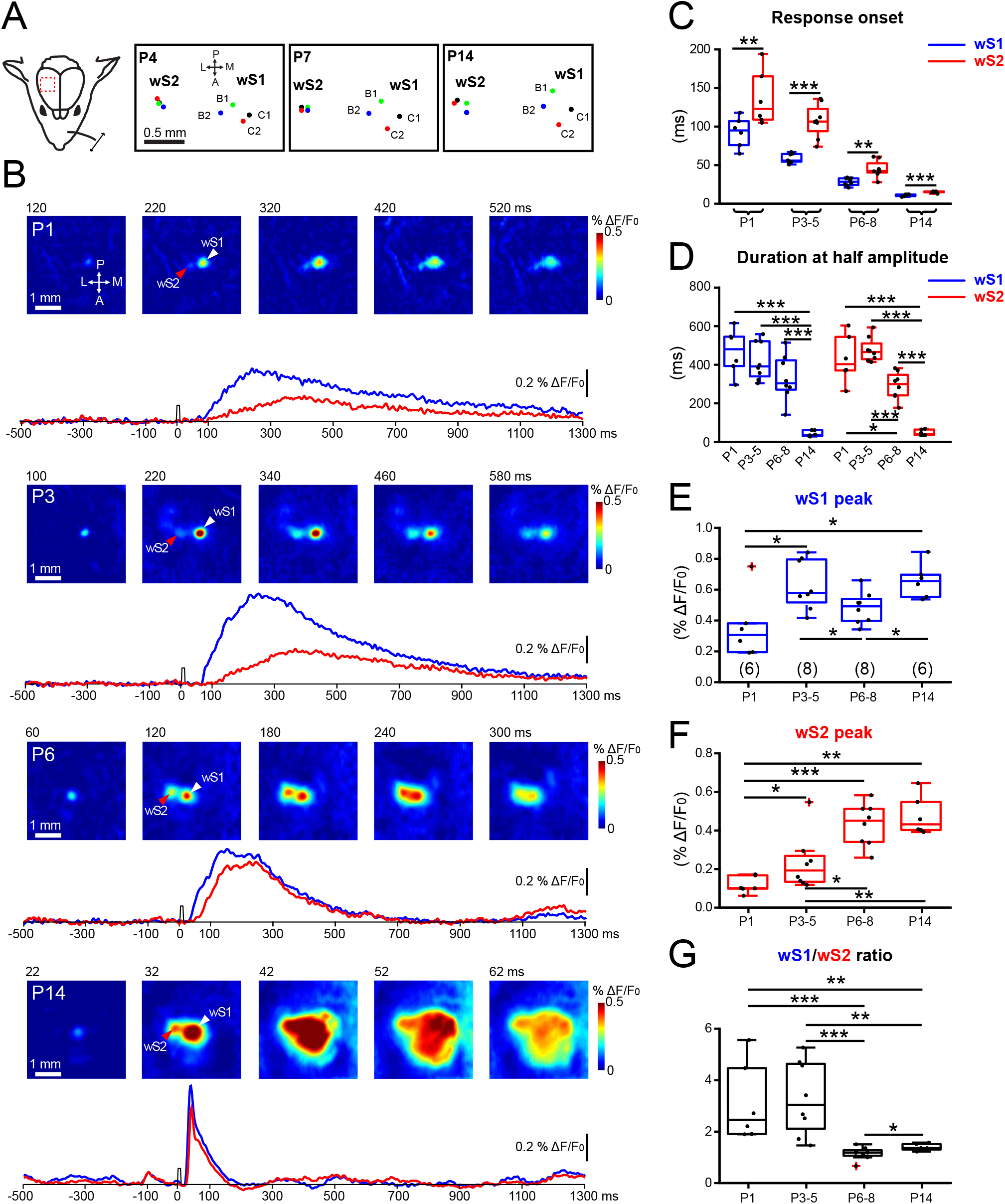
Developmental changes of sensory-evoked VSDI responses in wS1 and wS2. **(A)** Left: Schematic illustration of the experimental design. A single whisker deflection is performed together with VSDI of the whisker somatosensory cortex. Right: The spatial location of the representation of activity for four whiskers (B1, B2, C1, and C2) in wS1 and wS2 in a P4, P7 and P14 mouse is depicted. **(B)** VSDI of responses elicited by B3-whisker deflection in different age groups of WT mice (P1, P3, P6, and P14). The white arrowhead indicates the center of the wS1 response and the red arrowhead indicates the center of the wS2 response. The lower row shows 1.8-s long optical recording traces obtained by analyzing the signal in the centers of wS1 and wS2 over time (blue trace for wS1 and red trace for wS2). Time point of whisker stimulation at 0 ms. **(C)** The onset times of evoked responses in wS1 and wS2 in the different age groups. **(D)** The duration at the half-maximal amplitude of VSDI for different age groups in wS1 and wS2. **(E)** Box plot showing the evoked peak amplitude in the center of wS1 in the different age groups (n=6 barrels from N=3 P1 mice; n=8 barrels from N=4 P3-P4 mice; n=8 barrels from N=4 P6-P8 mice; n=6 barrels from N=3 P14 mice). **(F)** Box plot showing the evoked peak amplitude in the center of wS2 in the different age groups. **(G)** Box plot showing the evoked peak amplitude in wS1 divided by the evoked peak amplitude in wS2 in the different age groups. In box plots, black dots are the data points and the red + signs present outliers. In C, the stars indicate significant difference between wS1 and wS2 using a paired T-test. In D, the stars indicate significant difference between different age groups of wS1 and wS2 respectively, using an ANOVA with Bonferroni’s multiple comparison test. In E, F and G, the stars indicate significant difference between four age groups using the Mann-Whitney test. One star (*) indicates p<0.05, two stars (**) indicate p<0.01, and three stars (***) indicate p<0.001.

With VSDI a complex signal is recorded, which mostly arises from membrane fluctuations of the cells due to synaptic activity (Chen, Palmer, and Seidemann 2012). We first measured the duration at half-maximal amplitude for the two regions over time, which showed that it decreases in wS1 during the second postnatal week, while in wS2 already becomes shorter at P6-8 (Fig. 1D). This result may indicate that wS2 begins to receive faster inputs at the end of the first postnatal week, which would suggest a faster maturation of the inputs it receives at this time point compared to wS1. To assess the developmental time course of the strength of the inputs in the two cortical areas driven by whisker stimulation, we plotted the ΔF/F as a function of age for both regions independently. The data revealed that the overall activity generated in wS1 appears to reach a steady-state as early as P3, even though there is a decline at P6-8 with the signal going up again by P14 (Fig. 1E). In contrast, the maturation of incoming activity in wS2, compared to wS1, is delayed and reaches an upper plateau at P6-8 (Fig. 1F). These results suggest that inputs to wS1 develop first and wS2 follows. When plotting the ratio of activity between the two areas (wS1/Sw2) across age we find that the ratio approximates unity at P6-8 and only shows a marginal increase by P14 (Fig. 1G). This finding suggests that the amount of input-dependent excitation of wS1 and wS2 and the interaction between the two areas reaches a near stable state at the end of the first postnatal week.

### The spatio-temporal development of sensory-evoked output activity of wS1 and wS2 follows a bell-shaped curve

Having obtained a time-course of synaptic input development across the two postnatal weeks for wS1 and wS2, we next sought to assess how the inputs may translate to an action potential dependent output. We therefore applied transcranial calcium imaging recording from mice expressing the genetically encoded calcium reporter (GCaMP6) in neurons under the Snap25 promoter (Snap25-GCaMP6 mice), an approach that would primarily report action potentials. This allowed us to address the whisker-driven evoked responses with a mesoscopic spatial and temporal resolution of 5-15 μm and 100-200 ms, respectively. Similarly to the VSDI data, after a single whisker deflection, the two activity spots in wS1 and wS2 were apparent in the first 200 ms of recording and merged over time (Fig. 2A). This experimental paradigm was performed in five age groups that spanned from the first postnatal day to 2-month-old mice (Fig. 2B-E). We found that the highest evoked calcium response was reached for both regions during the first postnatal week, but unlike the VSDI, the peak activity for both areas was reached maximally in the P6-8 age group (Fig. 2C, D). When assessing the wS1/wS2 ratio of activity with age, we found that it started high in P0-2 (5.36 ± 0.8, n=4) and P3-5 (4.03 ± 0.5, n=11), and only started to decrease at the P6-8 time point (3.12 ± 0.23, n=16), after which it was gradually reduced at P14-16 (1.3 ± 0.04, n=12) and remained stable even at the P23-56 age group (1.41 ± 0.16, n=5). The wide-field calcium imaging results, together with the VSDI, reveal a window of increased input-output transformations taking place at the end of the first postnatal week for both wS1 and wS2, which may indicate the beginning of information transfer between the two cortical areas.

**Fig. 2.**
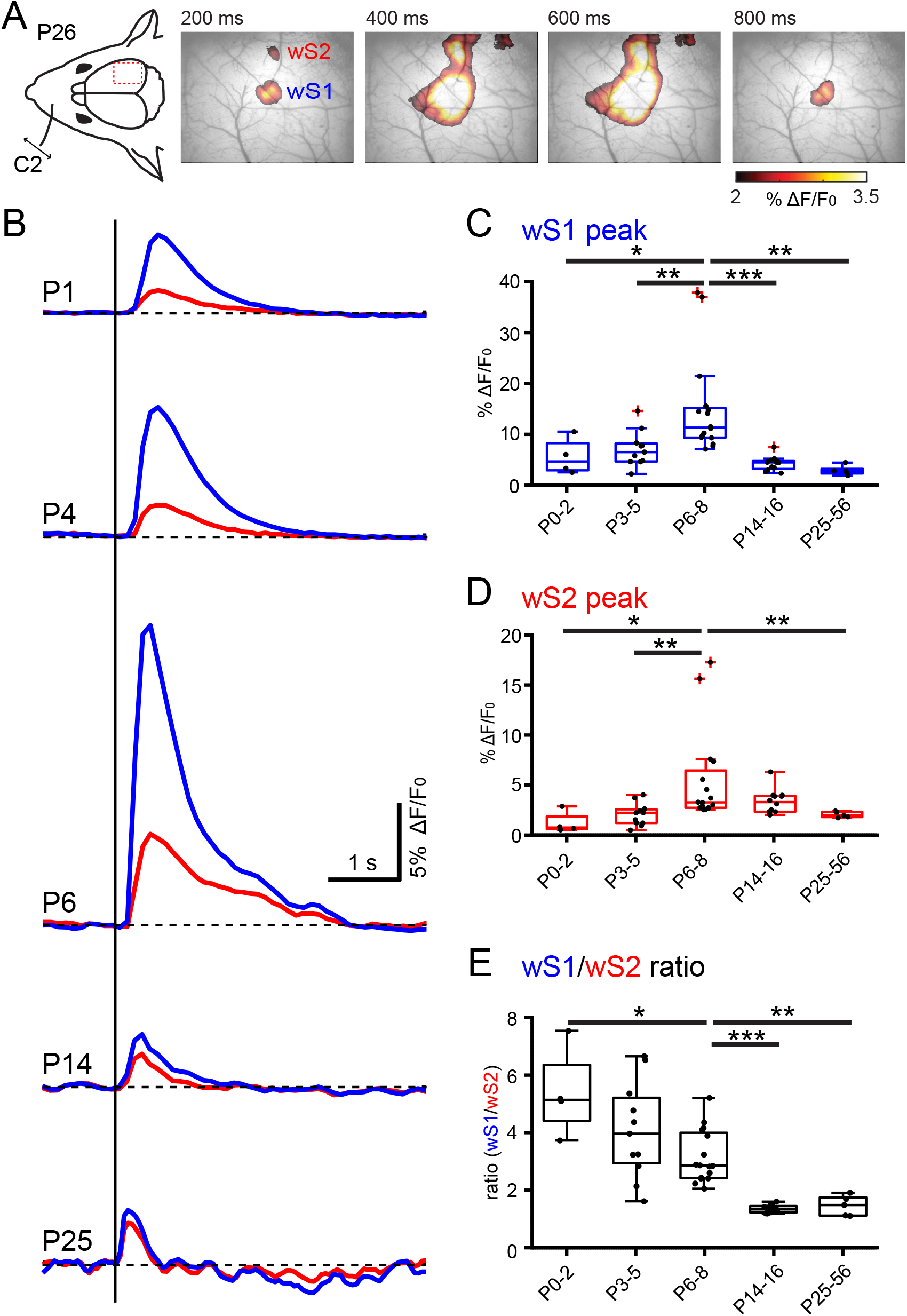
Developmental changes of sensory-evoked responses in wS1 and wS2 monitored with wide-field calcium imaging. **(A)** Left: Schematic illustration of the experimental design. A single whisker deflection is performed together with calcium imaging of the whisker somatosensory cortex of a P26 Snap25-GCamp6 mouse. Right: Evoked wide-field calcium imaging response at different time points after single C2 whisker deflection at 0 ms. Clearly separated responses in wS1 and wS2 are visible at 200 ms. **(B)** Temporal profile of wide field calcium imaging responses recorded in the center of wS1 (blue trace) and wS2 (red trace) at different ages (P1, P4, P6, P14, and P25 following C2-whisker stimulation. **(C)** Box plot displaying the evoked peak amplitude in the center of wS1 for different age groups (N=4 P0-P2 mice; N=11 P3-P5 mice; N=16 P6-P8 mice; N=11 P9-P12 mice; N=5 P25-P56 mice. **(D)** Box plot displaying the evoked peak amplitude in the center of wS2 for different age groups. **(E)** Box plot showing the evoked peak amplitude in wS1 divided by the evoked peak amplitude in wS2 for different age groups. In box plots, black dots are the data points and the red + signs present outliers.The stars indicate the significant difference between 5 age groups using the Mann-Whitney test. One star (*) indicates p<0.05, two stars (**) indicate p<0.01, and three stars (***) indicate p<0.001.

### The development of sensory-evoked spiking activity across layers in wS1 and wS2

To directly examine the action potential generation of each cortical layer and its precise time-course in wS1 and wS2 upon whisker stimulation, we performed acute *in vivo* silicon probe recordings at the four key age groups identified with wide-field imaging (P3-P5, P6-8, P14-P16 and P25-56). We performed the same whisker deflection paradigm and recorded spiking activity across all layers in wS1 and wS2 simultaneously. In order to accurately locate the probe insertion region for wS2, the Snap25-GCaMP6 mice were used, which allowed the identification of both areas online and in relation to the blood vessel pattern. Subsequently, an 8×8 silicon probe array was inserted in such an orientation to record activity in both wS1 and wS2 at the same time (Fig. 3A). By analyzing the local field potential (LFP) and calculating the Current Source Density (CSD) profile we were able to localize the recording sites along with the probes in respect to the cortical layers (Fig. 3B). We then extracted the Multi-Unit Activity (MUA) in the different layers for both regions across age. These analyses revealed significant developmental differences in the sensory activation of wS1 and wS2. The sensory evoked MUA pattern was rather short at P3-5, increased significantly at P6-8 and was short again in older age groups (Fig. 3C). These results are in line with the calcium wide-field imaging data, which also showed a developmental up- and down-regulation of the output in wS1 and wS2 centered at the end of the first postnatal week (P6-8). The analysis also showed that at the end of the second postnatal week, layer 4 (L4) of wS2 displays a sharp decline of spiking activity, indicative of the emergence of feed-forward inhibitory control, potentially coming from the thalamic inputs driving wS2 L4, or even cortical input coming from wS1 which would developmentally engage more wS2 L4 through long-range connections (Fig. 3D) (De León Reyes et al. 2019). This result is unexpected since it suggests that the higher-order area wS2 is regulated via developmental inhibitory control before primary wS1. The silicon probe data we obtained are in line with the calcium imaging results. The increased amplitude of the calcium imaging we observed at P6-P8 is underlined by a more prolonged spiking activity of the cortical circuit. In addition, this data also showed that the spiking activity in wS1 precedes that in wS2 in all age groups (Fig. 3E). Overall, these results demonstrate that wS2 is already strongly activated by sensory stimuli in the first postnatal week and undergoes more refined regulation during the second postnatal week.

**Fig. 3.**
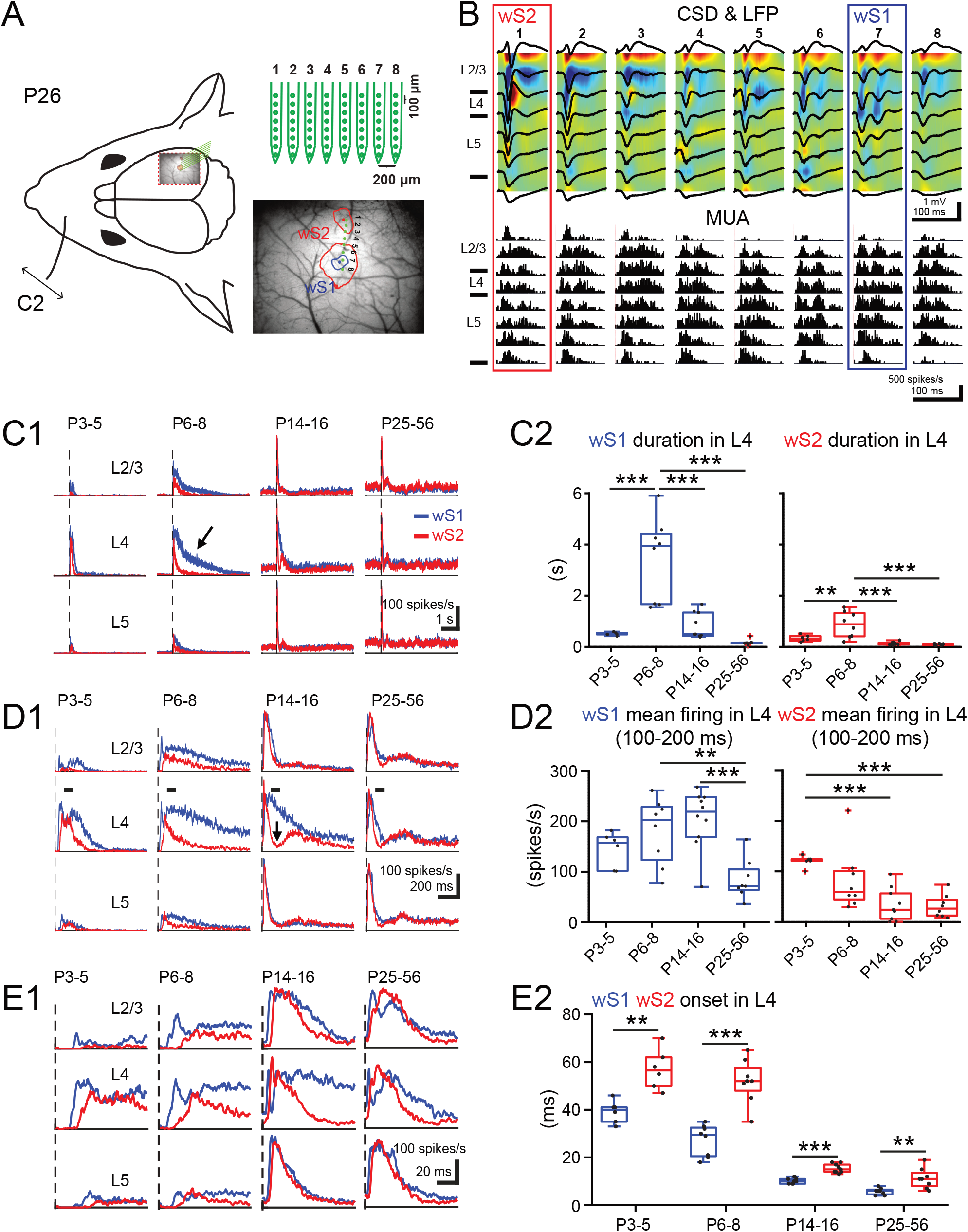
Developmental changes of sensory-evoked multi-unit-activity in wS1 and wS2. **(A)** Left: Schematic illustration of the experimental design showing the insertion position of an 8×8 multi-electrode probe array into wS1 and wS2 after identifying the center of evoked wS1 and wS2 by a single C2 whisker deflection in a Snap25-GCamp6 P26 mouse. Right: An example showing the evoked wide-field calcium imaging response (mean response from 11 to 40 ms after the onset of stimulation) in wS1 and wS2 by single C2 whisker deflection in a P26 mouse. We set 4% ΔF/F_0_ as the threshold to separate the evoked area of wS1 and wS2 (the red contour plot) and 5.5% ΔF/F_0_ as the threshold to confine the evoked area of wS1 (the blue contour plot). The blue dot indicates the center of evoked wS1 and the red dot indicates the center of evoked S2. The eight green dots indicate the insertion location of the 8×8 multi-electrode probe array. **(B)** Example of evoked local field potential (LFP) responses (black lines), color-coded current source density (CSD) plots and corresponding multi-unit activity (MUA) responses elicited by a single C2 whisker deflection in a P26 mouse. The cortical layers, L2-3, L4, and L5 were identified by the evoked CSD pattern. **(C1)** Grand averages of evoked MUA responses recorded in L2-3, L4, and L5 of wS1 and wS2 (1 s before and 5 s after onset of stimulation) in 4 age groups (n= 6 in P3-5, n= 8 in P6-8, n=10 in P14-16, n=8 in P25-56). **(C2)** Box plots showing the duration of evoked MUA of wS1 and wS2 in L4 of the different age groups. **(D1)** Same as in C1, but with higher temporal resolution showing only the first second after the onset of stimulation. **(D2)** The box plots show the mean firing rate of L4 in wS1 and wS2 between 100 to 200 ms poststimulus in the different age groups. **(E1)** Same as in C1, but with higher temporal resolution showing only the first 100 ms poststimulus. **(E2)** The box plots display the evoked onset time of MUA in L4 of wS1 and wS2 for the different age groups. In box plots, black dots are the data points and the red + signs present outliers. In C and D, the stars indicate a significant difference between 4 age groups using the ANOVA test. In E, the stars indicate the significant difference between wS1 and wS2 using a paired-T-test. One star (*) indicates p<0.05, two stars (**) indicate p<0.01, and three stars (***) indicate p<0.001.

### Postnatal genetic disruption of thalamocortical inputs arrests the developmental progression of wS2 sensory-evoked activity

In order to assess the dependence of wS2 whisker-driven activity on direct thalamocortical inputs versus via wS1, we sought to disrupt thalamocortical inputs at the level of the cortex, but not the thalamus. To achieve this, we used a genetically modified mouse line (Lhx2 conditional KO-cKO) in which the transcription factor *Lhx2* is floxed (Lhx2^f/f^) and postmitotically removed from cortical excitatory cells of all layers using the Nex-Cre mouse line (Goebbels et al. 2006). The loss of Lhx2 specifically in cortical neurons was shown to disrupt thalamic projections to the somatosensory cortex in the first postnatal week, barrel malformation, as well as reduced whisker-evoked activation of wS1 (Shetty et al. 2013; Wang et al. 2017). To assess when and how thalamocortical inputs may differentially affect the activation of wS1 and wS2 after whisker deflection, we performed concomitant VSDI in wS1 and wS2 in the cKO and WT for the Lhx2 allele mice. These experiments were performed in two age groups, P3-5 and P6-8. This choice is based on our previous functional experiments, which suggested that these age groups would represent direct thalamocortical activation only (P3-5), in addition to the start of wS1-wS2 communication (P6-8). The results showed that disrupting thalamocortical inputs to the somatosensory cortex in general, reduces the sensory-evoked activity in wS1 and wS2, albeit in a differential manner. At P3-5 there is a marginally stronger reduction in wS1 activation compared to wS2, whereas at P6-8 the effect is much stronger on wS2 (Fig. 4A-D). This is also reflected in the ratio of activation of wS1/wS2, which is not statistically different in the cKO mice compared to their littermate controls at P3-5, but it becomes significantly higher in cKO at P6-8 (1.16 ± 0.09 for WT vs. 2.33 ± 0.17 for KO, p<0.0001, Fig. 4E). This finding indicates that sensory information reaches wS2 mainly, if not exclusively, via the thalamus at P3-5. In contrast, by P6-8 the activation of wS2 after whisker deflection is a summation of both direct thalamic input and excitation coming from wS1, hence the stronger relative reduction of the overall activity of wS2 in the later age group.

**Fig. 4.**
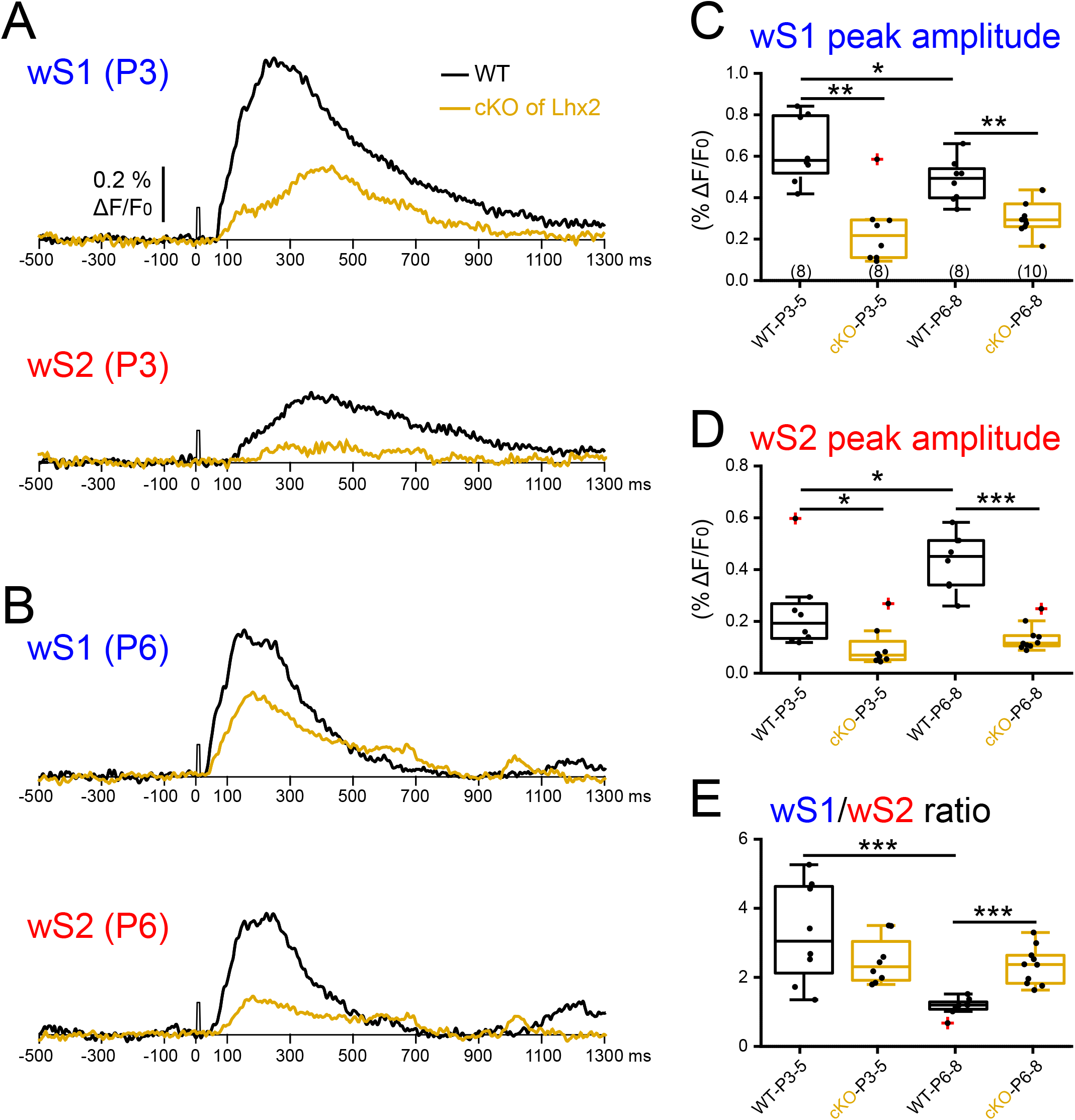
Postnatal removal of *Lhx2* from cortical excitatory cells leads to an arrest in the developmental progression of wS2 sensory-evoked activity. **(A)** An example trace of evoked VSDI recorded in the center of wS1 and wS2 from a P3 WT (black trace) and a P3 Lhx2 cKO mouse (orange trace). **(B)** An example trace of evoked VSDI recorded in the center of wS1 and wS2 from a P6 WT and a P6 Lhx2 cKO mice. **(C)** The box plots show the peak amplitude of evoked VSDI in wS1 of WT and cKO mice for the two age groups (n=8 recordings from N=4 P3-4 WT mice; n=8 recordings from N=4 P3-P4 cKO mice; n=8 recordings from N=4 WT P6-P8 mice; n=10 recordings from N=6 P6-8 cKO mice). **(D)** Same as in C, but for wS2. **(E)** The box plots show the evoked peak amplitude of VSDI in wS1 divided by the evoked peak VSDI amplitude in wS2 for the different age groups. In box plots, black dots are the data points and the red + signs present outliers. The stars indicate the significant difference between WT and cKO using the Mann-Whitney test. One star (*) indicates p<0.05, two stars (**) indicate p<0.01, and three stars (***) indicate p<0.001.

### Acute blockade of wS1 activity identifies a developmental window for information transfer between wS1 and wS2

It is known that in the adult, wS1 sends projections to wS2 and vice versa (El-Boustani et al. 2020). To assess if the changes that we observe in wS1 and wS2 VSDI between P3-5 and P6-8 are directly driven by the propagation activity from the former to the latter, we performed another set of silicon probe experiments where we acutely silenced action potential firing in wS1 and assessed spiking changes in wS2 after whisker deflection. We first tested if the application of tetrodotoxin (TTX) was able to silence the underlying activity in wS1 *in vivo*. Indeed, when recording spontaneous spiking in wS1, as well as whisker-evoked activity, we saw no action potentials occurring in the presence of TTX, indicating that this manipulation was efficient in abolishing activity for all the age groups we focused on, P3-5, P6-8, P14-16 and P26-46 (Fig. 5A). In contrast, analysis of the activity in wS2 after silencing wS1 showed no apparent effect in wS2 at P3-5. Nevertheless, there was a significant decline of wS2 spiking activity in the P6-8 age group in L2/3 and L4 (Fig. 5B), when analyzing the 100-1000 ms time window after the onset of whisker stimulation. The reduction of spike activity was also observed at P14-16, but it was only present in L2/3 and in the 50-80 ms time window after the onset of whisker stimulation (Suppl. Fig. 1). In adolescent/adult animals, although abolishing activity in wS1 leads to no observable difference in wS2 with single time deflection, an increased wS2 activity was observed upon repetitive whisker stimulation at 10 Hz (Suppl. Fig. 2). This result suggests the maturation of feed-forward inhibition from wS1 to superficial layers of wS2 after the end of the second postnatal week. These findings support the calcium imaging and VSDI data in control and cKO mice and point at a developmental window for excitatory information transfer between wS1 and wS2 at the end of the first and beginning of the second postnatal week, indicating the emergence of higher-order cortical processing of sensory information. The addition of inhibitory control in the transfer of activity from wS1 to wS2 in the third postnatal week coincides with the onset of active exploratory whisking behavior and most likely represents an extra level of regulation of the information necessary to build proper sensory representations in wS2.

**Fig. 5.**
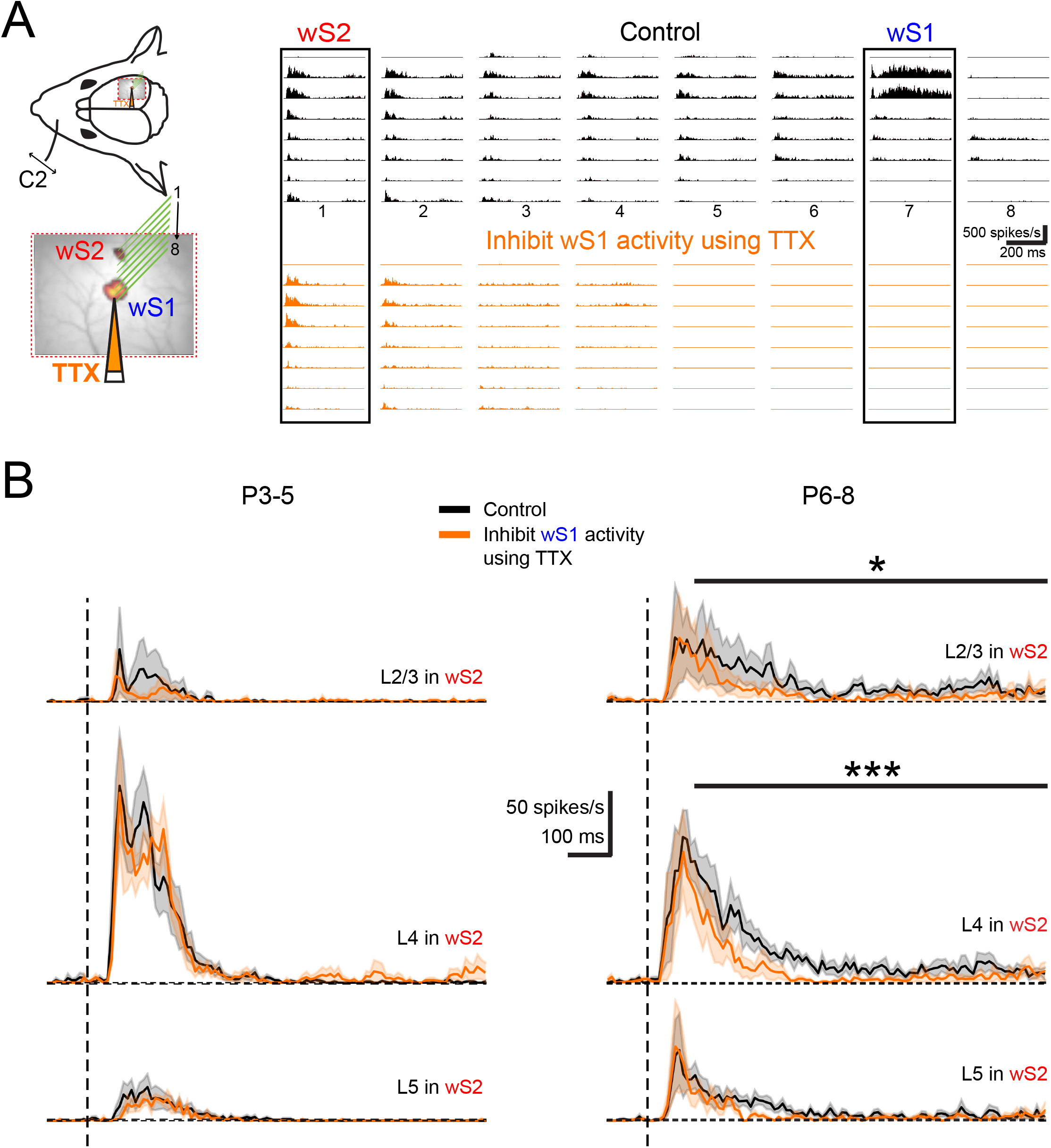
Acute Inhibition of wS1 activity differentially influences sensory-evoked responses of S2 over development. **(A)** Left: A schematic illustration showing the insertion position of an 8×8 silicon probe array in wS1 and wS2. TTX was injected into wS1 through a micro-glass pipette (orange). Right: An example of evoked MUA recorded in wS1 and wS2 after C2 whisker deflection in a P826 mouse in the control condition (black), and after local TTX injection in wS1 (orange). **(B)** Left: Average traces of evoked MUA in L2-3, L4, and L5 of the P3-5 age group (n=5) before (black) and aer TTX injection in wS1 (orange). Right: Average traces of evoked MUA in L2-3, L4, and L5 of P6-8 age group mice (n=6) before (black) and after TTX injection in wS1 (orange). A statistical comparison was performed between control and TTX injection conditions using the paired T-test for the me window of 100 to 1000 ms after the onset of whisker smulation. One star (*) indicates p<0.05 and three stars (***) indicate p<0.001.

Overall, our data identify a developmental time window in which activity from wS1 starts impacting that of wS2 upon the presentation of a sensory stimulus. Even though prior to P5 both wS1 and wS2 already receive functional thalamocortical inputs, there is little communication between the two cortical areas (Suppl. Fig. 3). It is only at the end of the first postnatal week that wS2 starts to receive functional inputs from wS1, and this communication becomes more refined during the second postnatal week with the appearance of feed-forward inhibition in L4 between the two areas (Suppl. Fig. 3).

## 3. Discussion

The development of cortico-cortical communication in mammals is fundamental for the emergence of higher-order computations. These computations include the processing of sensory information by a number of cortical regions to extract the fundamental features of a stimulus within a given context. The sequential engagement of cortical regions in processing sensory stimuli has its prototypical example in the visual system and the two identified streams of information flow in primates and potentially rodents (Glickfeld and Olsen 2017; Stone, Dreher, and Leventhal 1979). In the rodent whisker-related somatosensory cortex, this process involves not only the activation of the primary cortical region, called the barrel cortex (wS1), but also of the secondary region (wS2), two areas which have been found to be activated upon a tactile event received by the whiskers (Condylis et al. 2020; Feldmeyer et al. 2013; Hubatz et al. 2020). Even though wS2 receives direct inputs from the thalamus, specifically the POm and the ventral lateral part of the VPM (VPMvl) (Chakrabarti and Alloway 2006; El-Boustani et al. 2020), which can provide its direct sensory stimulus-driven activation, it also receives connections from wS1. A number of studies have examined the response properties of neurons in wS2 after whisker deflection and have found that they have larger receptive fields than neurons in wS1 (Goldin et al. 2018; Kwegyir-Afful and Keller 2004). This also matches the less segregated representation of individual whiskers seen with VSDI, suggesting the integration of individual whisker responses from wS1 to wS2 (Hubatz et al. 2020). In this study, we examined when and how wS1 begins communicating with wS2 over the course of postnatal development, in order to assess when higher-order processing of whisker stimuli commences.

By utilizing two wide-field imaging approaches, one of which provides a readout of the overall synaptic inputs a region receives (VSDI), whereas the other provides a readout of spiking (calcium imaging), we were able to identify the time course of input-output transformations in wS1 and wS2. VSDI showed that wS1 inputs from the thalamus develop a few days before the ones in wS2, and are already present within the first two days after birth. In contrast, utilizing wide-field calcium imaging from all neurons in the two regions, again simultaneously, revealed that their output is more synchronized and tracks with the increased inputs that wS1 receives. Our follow-up silicon probe-based electrophysiological recordings in wS1 and wS2 consolidated the calcium imaging data and provided insights into the temporal domain of sensory evoked spiking activity across cortical layers. The results show that during the first postnatal week, whisker deflection leads to a faster onset of wS1 spiking activity compared to wS2, a difference that gets reduced with time and reaches its lowest point in adult mice. The recordings also reveal that there is an increase in the sustained spiking activity of the two regions at P6-8, albeit more so in the wS1 compared to wS2. This time window coincides well with the increase in the generation of lateral excitatory connectivity between pyramidal cells of L2/3 in wS1 (Bureau, Shepherd, and Svoboda 2004), thereby providing an underlying circuit-based explanation for the large increase in sustained activity in wS1, which is transferred to wS2 after a whisker stimulus. Surprisingly, we also found that the sustained excitability of layer 4 in wS2 at P6-8 is curtailed at P14-16, much more strongly than in wS1. This finding would suggest that a type of sensory-driven feed-forward inhibitory (FFI) control of wS2 develops earlier than in wS1, the latter only showing the same pattern in layer 4 after P26. As a higher-order cortical area, one would have expected that FFI in wS2 would lag behind wS1 in the addition of such a control mechanism. This is not what we nevertheless observe, with our findings suggesting the earlier maturation of certain aspects of signal processing in wS2 compared to wS1.

In order to assess the direct sensory-driven thalamocortical activation of wS1 and wS2 versus through each other, we took advantage of a genetically modified mouse line that displays disrupted thalamocortical projections upon removal of the key transcription factor *Lhx2* from cortical cells. Since the disruption of the thalamocortical inputs in this mouse line is expected to be of the same magnitude for both cortical areas, any difference we would observe in their activation ratio after whisker stimulation would be attributed to defects in additional activity coming from other cortical areas. Indeed, we observed a much more pronounced reduction of wS2 activity compared to wS1 at P6-8 in cKO mice, but not at P3-4. This suggests that initially the thalamocortical inputs in the two areas are equally disrupted, whereas a few days later, wS2 has also “lost” the indirect sensory-driven activity that drives it further via wS1. Based on these findings, we directly tested the contribution of wS1 activity to sensory-driven activation of wS2 via inhibition of action potential firing in wS1. In line with the cKO, we find that this acute activity manipulation at P6-8 reduces the activity in all layers of wS2, with the larger difference observed in layer 4 (L4). The layer-dependency of this effect matches to a large extent the anatomical axonal innervation that has been reported in adult mice between wS1 to wS2, appearing in more or less all layers of wS2 and also heavily innervating L4 (Minamisawa et al. 2018). Interestingly the innervation between L4 cells of the two areas may be even more pronounced at P6-8, since it has been observed that L4 wS1 axons from stellate cells send long-range projections contralaterally that subsequently trim before the end of the second postnatal week (De León Reyes et al. 2019). This phenomenon would also leave open the possibility that there are more abundant long-range connections between L4 cells of wS1 and wS2.

In line with the wide-field results, prior to P6-P8, we observed no change in the spiking of wS2 after acutely blocking activity in wS1. A few days later, at the end of the first postnatal week (P6-P8) we saw that information is fed-forward from wS1 to wS2, as indicated by the reduction of the late activation of the latter, when abolishing activity in the former. A week later (P14-16) and at the start of active whisking behavior, blocking wS1 activity decreases the spiking activity 50-80 ms in L2/3 followed by a negligible increase of spiking. This bivalent effect in the early versus late activity at P14-16 seems to suggest that the period between the second and third postnatal week is a transition one, since after that (>P26), this effect is not observed. In contrast, an overall inhibitory influence of wS1 onto wS2 was pronounced upon repetitive deflection of the whiskers at 10 Hz, which is in line with a known temporal bias in information transfer to wS2 (Melzer et al. 2006) and matches the whisking frequency of adult animals (De León Reyes et al. 2019). This finding in adult animals is nevertheless intriguing, as it is generally considered that wS1 would transmit information to wS2 in a positive feed-forward sequential manner. Even though this is not what we find, it may well be the case when the animal is performing a behavioral tactile-dependent task (Chen et al. 2015), despite the fact that to our knowledge inactivating wS1 and assessing the activity in wS2 has never been directly tested in rodents. Nevertheless, there is evidence from experiments in cats, rabbits and marmosets that support a parallel processing model of information coming directly from the thalamus, compared to a sequential wS1 to wS2 one (Burton and Robinson 1987; Manzoni et al. 1979; Turman et al. 1995; Zhang et al. 1996). These studies have shown that a large number of neurons in wS2 show no change in their sensory-driven activation when wS1 was acutely, and in some cases reversibly, inactivated, whereas others display a small decrease. Based on these findings, and recent ones showing the activation of wS2-projecting wS1 neurons upon a whisker touch and the acquisition of texture discrimination, more functional experiments are required in awake behaving mice to determine the direct impact of wS1 activity to wS2 in adulthood. Interestingly, published work has provided some insights into the development of parallel versus sequential activation between somatosensory paw area S1 (pS1) and paw primary motor cortex (pM1) of rat (Gómez et al. 2021). The work proposes that at P8, paw sensory stimulation reaches pS1 and pM1 through parallel streams, potentially from the thalamus, whereas at P12 it reaches pM1 via pS1 through a serial activation scheme.

Overall, our data provide novel insights into the developmental transfer of information between primary and secondary somatosensory areas. We have revealed a previously unidentified developmental time window for the positive information flow between wS1 and wS2 at the end of the first postnatal week, which suggests the beginning of more complex representations of the sensory environment in the cortex.

## 4. Methods

### Animals

Animal experiments were approved by the Cantonal Veterinary Office Zurich, the local German ethics committee (#23177-07/G10-1-010), and the Academia Sinica Institutional Animal Care and Use Committee in Taiwan. Animals were in husbandry with a 12-h reverse darklight cycle (7 a.m. to 7 p.m. dark) at 24 °C and variable humidity. VSD imaging was performed on C57BL6J mice. *Lhx2* conditional knockout (cKO, *Lhx2*^f/f^; *Nex*-Cre) mice were generated as described (Chou et al. 2009; Goebbels et al. 2006). All lines were maintained on a C57BL/6 background. Wide field imaging and multi-electrode recordings were performed on Snap25-2A-GCaMP6s-D mice.

### Animal surgery

We used 77 Snap25-GCaMP6s mice at the ages from P0 to P56 for wide field imaging and multi-electrode recordings. Mice were anesthetized by urethane throughout the whole experiment. A heating pad was used to maintain the mouse body temperature at 37°C. The depth of anesthesia was checked with breathing speed and paw reflexes throughout the experiment.

The skull of the right hemisphere was exposed by removing the skin on top, and a metallic head holder was implanted on the skull with cyanoacrylate glue and dental cement. 20G needle was used to open ∼3mm*3mm cranial window which exposed both S1 and S2. Extreme care was taken not to cause damage or surface bleeding in neonatal pups.

### Whisker stimulation

A single whisker was stimulated 1 mm from the snout in rostral-to caudal direction (about 1 mm displacement) using a stainless-steel rod (1 mm diameter) connected to a miniature solenoid actuator. The movement of the tip of the stimulator bar was measured precisely using a laser micrometer (MX series, Metralight, CA, USA) with a 2500 Hz sampling rate. The stimulus takes 26 ms to reach the maximal 1 mm whisker displacement, with a total duration of 60 ms until it reaches baseline (Yang et al. 2017).

### VSD imaging

The procedure of VSDI was according to our previous report with small modifications (Luhmann 2016; Yang et al. 2013). Briefly, the VSD RH1691 or RH2080 (Optical Imaging, Rehovot, Israel) was dissolved at 1 mg/ml in Ringer solution containing (in mM): 135 NaCl, 5.4 KCl, 1 MgCl_2_, 1.8 CaCl_2_ and 5 HEPES (pH was set to 7.2 with NaOH). The VSD was topically applied to the surface of the opened skull and allowed to diffuse into the cortex for 40-60 min. Subsequently, the unbound dye was carefully washed away with Ringer solution. The cortex was covered with a 1% low-melting agarose and a coverslip was placed on top to stabilize the tissue. Imaging was performed using a MiCam Ultima or MiCAM05 high-speed camera (Scimedia, Costa Mesa, CA, USA) with 625 nm excitation and 660 nm long-pass filtered emission. The field of view was 3.1×3.1 mm^2^ with 100×100 pixels for recordings using the MiCam Ultima and 6.8×6.8 mm^2^ with 256×256 pixels for recordings using the MiCAM05.The frame sampling frequency was 500 Hz. The evoked activity following whisker stimulation was averaged from 5 recording sessions. Fluorescence signals were analyzed using custom-made routines in Matlab (Mathworks, Natick, MA, USA). The fluorescence change (ΔF/F_0_) was calculated as the change of fluorescence intensity (ΔF) in each pixel divided by the initial fluorescence intensity (F_0_) in the same pixel. The center of a single barrel was functionally determined by the local maxima of the initial appearance of the VSDI response (see Fig. 1A). The fluorescence signals in the functional center of the barrel were used to analyze the peak amplitude, and onset time. The onset time was detected by the threshold in 1-fold baseline SD.

### Wide field calcium imaging

The principal whisker-related S1 barrel columns (B1, B2, C1 and C2) and their corresponding representations in S2 were identified by wide field calcium imaging. Before P16, the cortical surface was visualized through the intact skull. In mice older than P21, the skull was thinned to have better visualization of calcium signal during wide field imaging. By shining blue light (488 nm LED) on the cortical surface, the functional maps in S1 and S2 were revealed by stimulating the principal whiskers. Images were acquired through a 1x Nikon objective or 1.25x Olympus objective with a CCD camera with 5 or 10 fps. The recordings lasted 10 s with a 2 s baseline and 8 s post-stimulation period. The wide field calcium signals obtained for principal barrels were then mapped to the blood vessel reference image and used to guide the location of the craniotomy and subsequent *in vivo* multi-electrode recording.

### In vivo multi-electrode recordings

S1 and S2 neural activities were recorded simultaneously with a 64-channel silicon probe inserted perpendicularly into the cortex. Each of the 8 shanks has 8 recording sites (100 µm apart). The distance between each shank is 200 µm (NeuroNexus Technologies, Ann Arbor, M). Insertion of the silicon probe was guided by the wide field calcium imaging results. A silver wire was placed into the cerebellum as a ground electrode. Before insertion, the silicon probe was dipped into DiI solution therefore the insertion points were marked after removing the probe. All data were acquired at 20 kHz and stored with MC_RACK software (Multi Channel systems). The total duration of multi-electrode recordings varied between 3 h and 5 h.

After a capillary containing 2 µM TTX was inserted into S1 without injecting TTX into the cortex, 20 trails of single and 10 Hz whisker stimulations were applied and S1 and S2 activities were recorded. Afterwards, 200 nl of TTX in the capillary was injected into the cortex to block activity in S1. The same stimulation paradigm was performed again and S1 and S2 activities were recorded.

### Analysis of wide field calcium imaging data

Wide field calcium imaging data were analyzed using a custom-made Matlab script (Matlab 2019a, Mathworks, MA, USA). The fluorescence change (ΔF/F_0_) was calculated as the change of fluorescence intensity (ΔF) in each pixel divided by the baseline fluorescence intensity (F_0_, average of 500 ms absolute fluorescence before the onset of whisker deflection) in the same pixel. The S1 and S2 evoked activity over different age group were compared by using the highest evoked calcium response.

### Analysis of *in vivo* multi-electrode silicon probe data

Extracellular silicon probe data were analyzed using a custom-made Matlab script (Matlab 2019a, Mathworks, MA, USA). The raw data signal was band-pass filtered (0.8-5 kHz) and the multi-unit activity (MUA) was extracted with the threshold of 5 times the standard deviation (SD) of baseline. The current source density (CSD) map was used to identify L2-3, L4 and L5. The earliest CSD sink was identified as layer 4, followed by L2-3 and L5 (van der Bourg et al. 2017; Reyes-Puerta et al. 2015). The MUA were separated and the average MUA were calculated and smoothed by 5 ms or 10 ms sliding window averaging for each stimulation paradigm. For S1 and S2 evoked activity onset and duration, a threshold of baseline +/- SD was set. For comparing S2 evoked activity in control vs TTX, the mean firing rate was calculated in two periods (0 to 100 ms and 100 to 1000 ms after onset of whisker deflection) for single whisker deflection. For whisker stimulation at 10 Hz, the mean firing rate was calculated in the time window after each whisker deflection.

### Statistical analysis

The statistics are indicated for every experiment in the manuscript or figure legend. The Mann-Whitney test, paired T-test and one-way-ANOVA were used to perform the statistical analyses.

## Acknowledgements

We would like to thank all members of the Karayannis lab for the comments and feedback on this project, as well as Leonardo Facheris for help with some of the analysis of the data.

## Competing interests

We declare no financial and non-financial competing interests on behalf of all authors.

---

**Suppl. Fig. 1.**
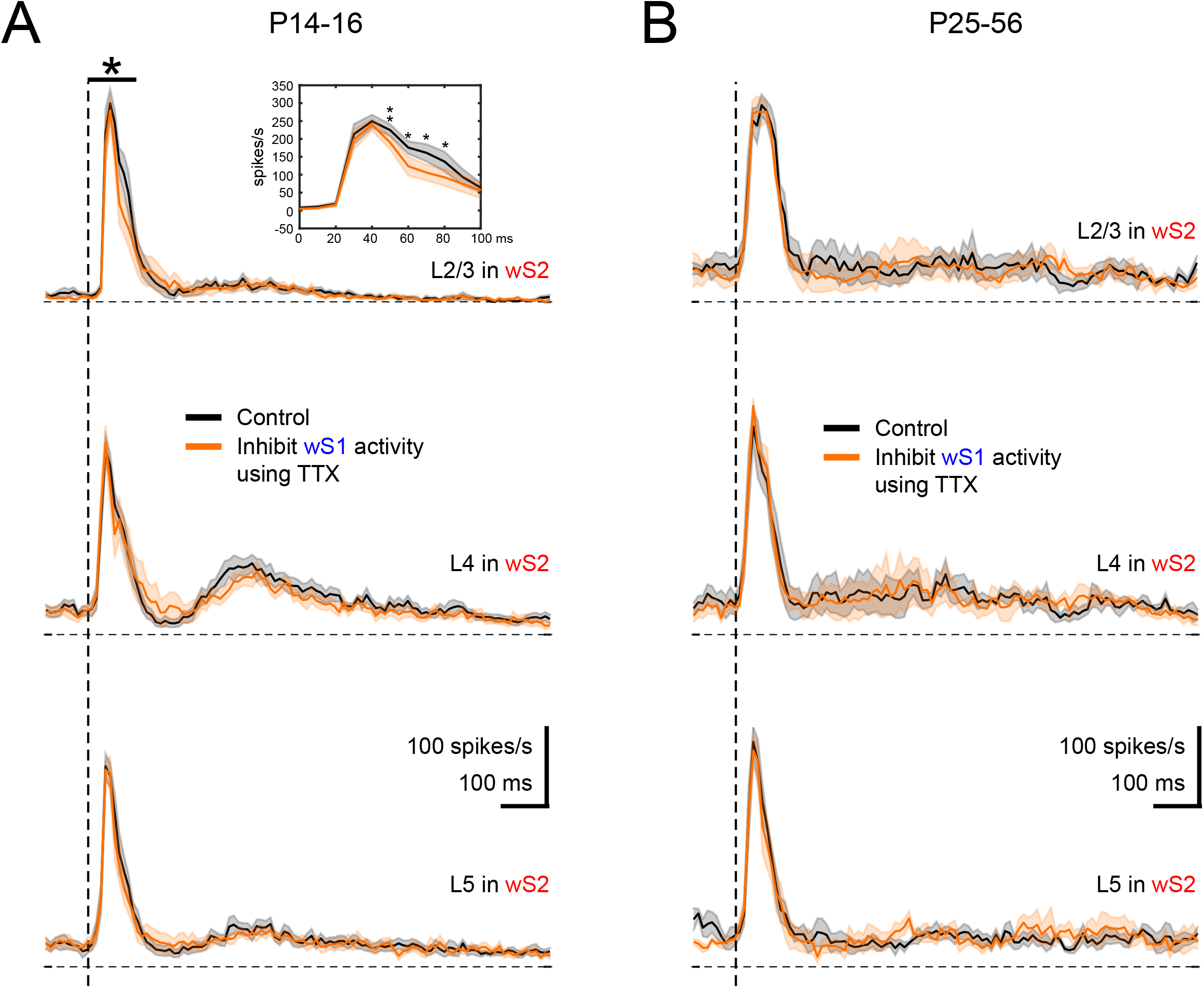
Inhibiting S1 influences the sensory-evoked response of S2 by single whisker deflection. **(A)** Average of evoked MUA response in L2-3, L4, and L5 of P14-16 age group mice (n=9) in the control condition (black) and TTX injection in S1 (orange). The star indicated p<0.05 using paired t-test during the period of 0 to 100 ms after the onset of whisker smulation. The insect shows the detailed paired t-test comparison of mean firing rate between control and TTX injection conditions using 10 ms bins during the period of 0 to 100 ms after the onset of whisker smulation. **(B)** Average of evoked MUA response in L2-3, L4, and L5 of P25-56 age group mice (n=5) in the control condition (black) and TTX injection in S1 (orange).

**Suppl. Fig. 2.**
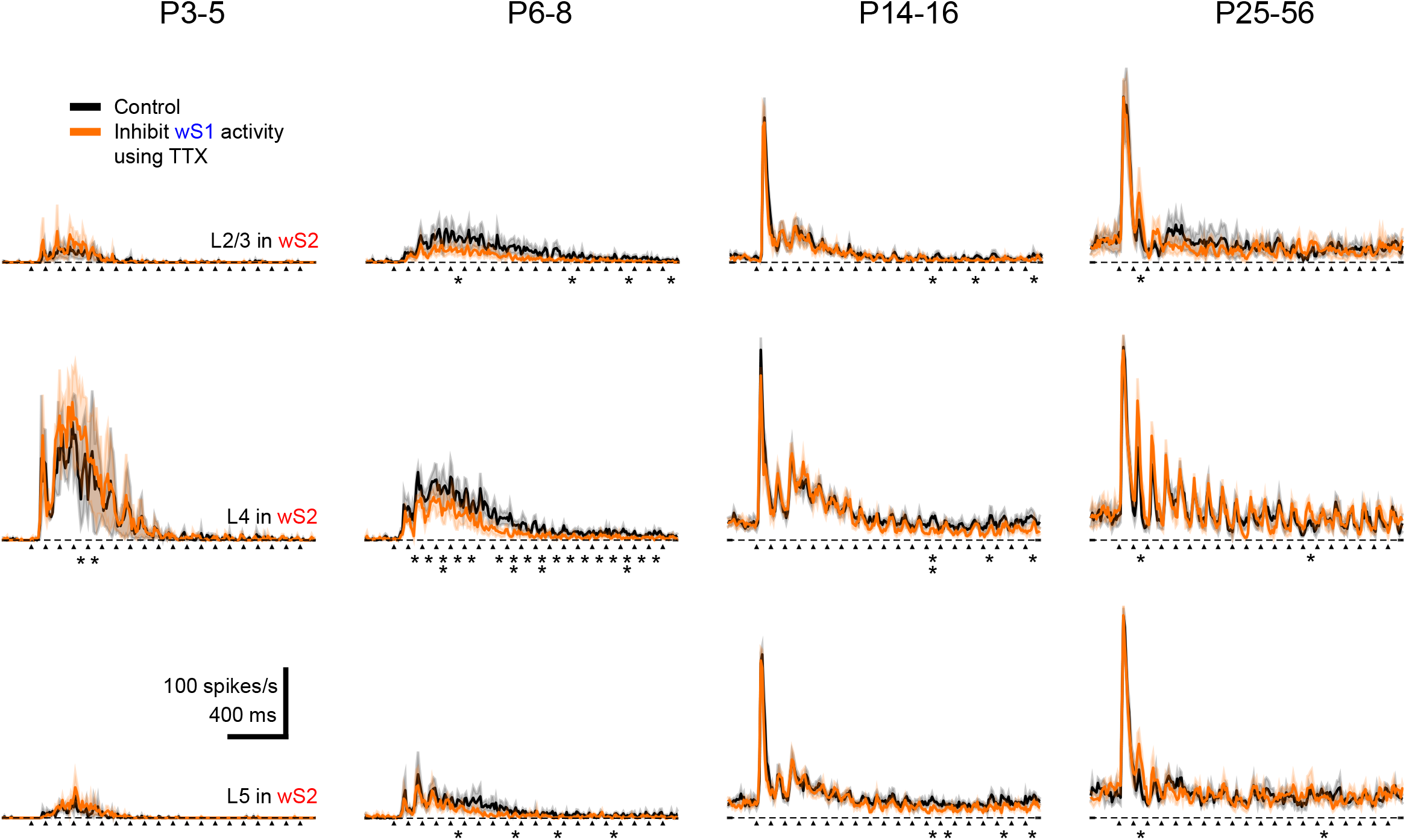
Inhibiting S1 influences the sensory-evoked response of S2 by single whisker deflection at 10Hz for 2sec. Average of evoked MUA response in L2-3, L4, and L5 of P3-5 (n=3), P6-8 (n=6), P14-16 (n=7), and P25-56 (n=5) age groups of mice. The arrows indicate the onset of whisker deflection. The star indicated the significant difference between control and TTX injection conditions using the paired t-test during the period of 0 to 100 ms after the onset of whisker stimulation. One star indicates p<0.05 and two stars indicate p<0.01.

**Suppl. Fig. 3.**
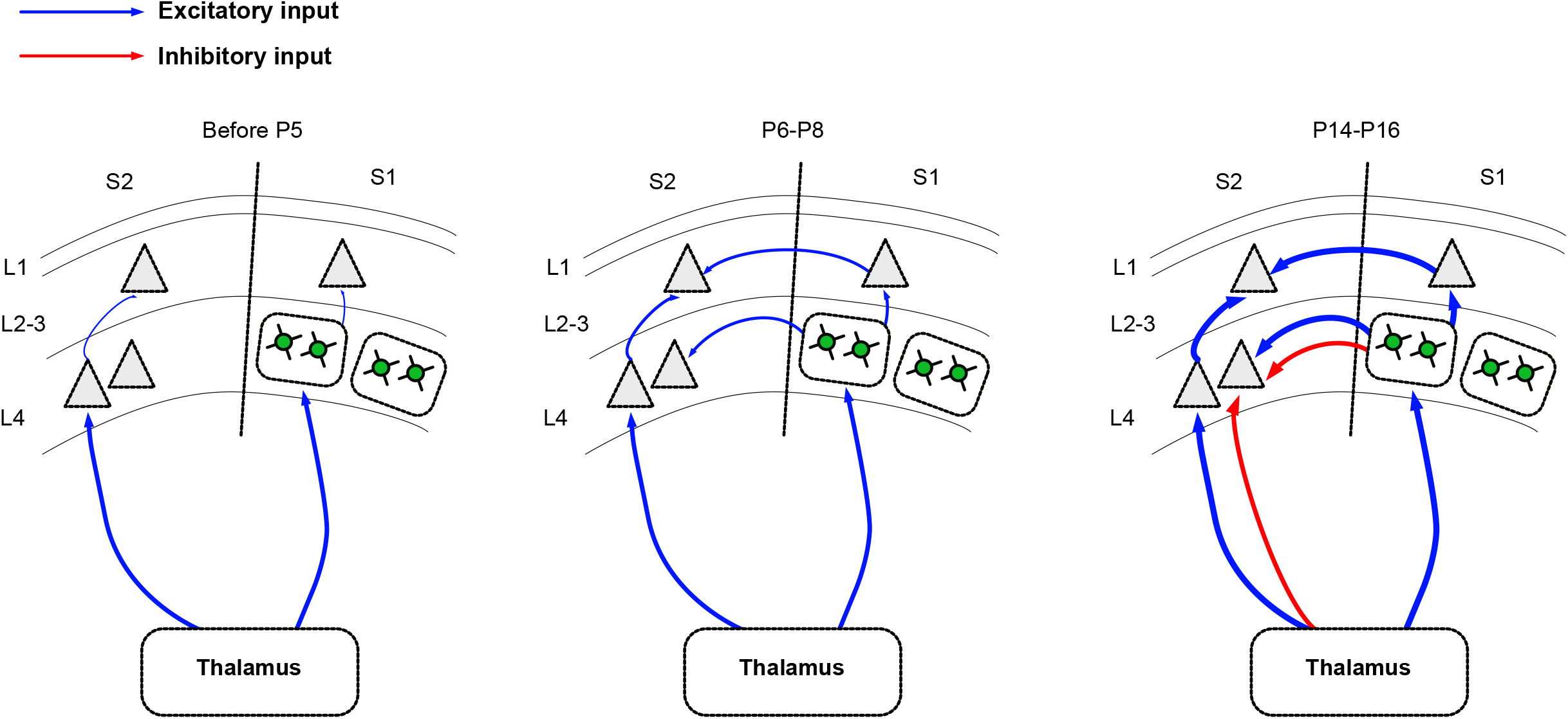
Schematic of the developmental functional connectivity between wS1 and wS2. Thalamocortical input: Both wS1 and wS2 already receive input from the thalamus shortly after birth and this input becomes stronger during the second postnatal week. Corctico-cortical connectivity: Before P5, L2-3 of wS1 and wS2 receive very little input from the respective L4 and functional connectivity from wS1 to wS2 is not yet developed. At P6-8, L2-3 in both wS1 and wS2 receive more activation upon whisker stimulation, and wS2 L2-3 and L4 start receiving inputs from wS1. At the end of the second postnatal week, wS2 starts receiving inhibitory regulation from either the thalamus or wS1 or both while the excitatory inputs mature.

## Notes

### Competing Interest Statement

The authors have declared no competing interest.

